# Structural studies of hepatitis C virus non-structural protein-5b of genotype 4a

**DOI:** 10.1101/2022.04.06.486972

**Authors:** Hanaa Gaber, Dierk Niessing, Ulrike Protzer, Robert Janowski

## Abstract

Hepatitis C virus genotype 4a non-structural protein-5b is an RNA-dependent RNA-polymerase responsible for the efficient virus genome replication. It has been of great interest as a drug target. The availability of the crystal structure of HCV genotype 4a NS5b would facilitate the structure-based drug design. Here we report the X-ray structure of NS5b solved at 3.1 Å resolution, setting the groundwork for the above-described therapeutic strategy.

**Synopsis:** X-ray structure of hepatitis C virus genotype 4a non-structural protein-5b, RNA-dependent RNA-polymerase responsible for the efficient virus genome replication.

## 1. Introduction

Determination of the three-dimensional (3D) structure on the atomic level provides scientists with deep insights into viruses’ world. The overall 3D structure reveals the virus protein shape and the arrangement of the active sites, which enables a structure-based understanding of the virus life cycle and the interactions between the virus and host molecules. Numerous methods are applied to resolve protein structure, including, X-ray crystallography, electron microscopy and NMR. The hepatitis C virus (HCV) genome vary between different genotypes by as much as 20-30%. Accordingly, more than 20,000 sequences covering different HCV genotypes are available. In addition, 3D structures of HCV proteins were elucidated for different genotypes (Qiu *et al*., 2015). A range of anti-HCV drugs has been developed targeting non-structural proteins, in particular, NS5b. The HCV NS5b protein is RNA-dependent RNA polymerase (RdRp), which replicates the viral RNA genome by using an intermediate negative-strand RNA template. Therefore, NS5b arose as a very important target for HCV inhibition. Recently, drugs targeting NS5b of different HCV genotypes were approved showing antiviral efficiency in up to 95% in native patients (Degasperi & Aghemo, 2014, Asselah & Marcellin, 2015, Bunchorntavakul & Reddy, 2015, Loustaud-Ratti *et al*., 2016, Nakamura *et al*., 2016). Approximately, there are over 200 crystal structures available for HCV NS5b covering 1a, 1b, 2a, and 2b genotypes, whereas no structure is elucidated for NS5b of genotype 4a. Of note, HCV genotype 4a accounts for 13% of the worldwide infections, with a majority of the load in the Middle East, Northern Africa and Sub-Saharan Africa including the highest infection rate of 12-15% in Egypt.

The aim of our study was to solve the structure of NS5b of genotype 4 HCV to allow for predictions whether NS5b inhibitors developed for HCV genotypes 1 to 3 would also bind the genotype 4 polymerase. Here, we report the X-ray structure of HCV NS5b of genotype 4a at a resolution of 3.1 A.

## 2. Materials and methods

### 2.1. Macromolecule production

For the PCR amplification of NS5b lacking 23 C-terminal amino acids, the ED43/SG-Feo (ED43; genotype 4a, kindly provided by Prof. Dr. Charles Rice) plasmid was used as a template together with HCV-GT4a-NS5b-XbaI-RBS (Fw) and HCV-GT4a-NS5b-XhoI (Rv) primers (Table 1). PCR product and pET-30b (+) (Novagen) were digested with XbaI and XhoI restriction enzymes, and then ligated using T4 DNA ligase. The ligated DNA vector, was transformed into *E. coli* BL21 (DE3) competent cells. To induce protein expression, isopropyl β–D–1–thiogalactopyranoside (IPTG, 0.5 mM) was added and cultured cells were cooled to 32°C for 4 hours. Cells were harvested by centrifugation at 10,000 x g and 4°C for 30 minutes and the pellet was stored at –80 °C until use. To confirm the expression of His-Tagged HCV NS5b, western blot analysis has been performed (Supplementary materials figure 1). The pellet from 3 L of culture was resuspended in buffer A (50 mM TRIS pH 7.5, 500 mM NaCl, 20 mM imidazole pH 7.5), supplemented with DNase and EDTA-free protease inhibitor cocktail to a total volume of 50 ml. After sonication (4 × 4 min, amplitude 30%, output: 6), the lysate was clarified by centrifugation (39000 x g, 30 min), loaded onto 5 ml HisTrap column (GE Healthcare), equilibrated in buffer A, and washed with 20 column volumes of buffer A. High salt wash was implemented (50 mM TRIS pH 7.5, 1 M NaCl, 20 mM imidazole pH 7.5). His tagged protein was eluted from the column in 10 CV linear imidazole gradient using buffer B (50 mM TRIS pH 7.5, 250 mM NaCl, 500 mM imidazole pH 7.5). Protein-containing fractions were pooled, concentrated and loaded onto a size exclusion chromatography column (Superdex 200, 16/60) equilibrated with SEC buffer (20 mM TRIS pH 7.5, 200 mM NaCl). The peak fractions were analysed in 12% SDS-PAGE. The selected fractions were then pooled and concentrated to 4 mg ml^-1^ by ultrafiltration.

**Table 1.**
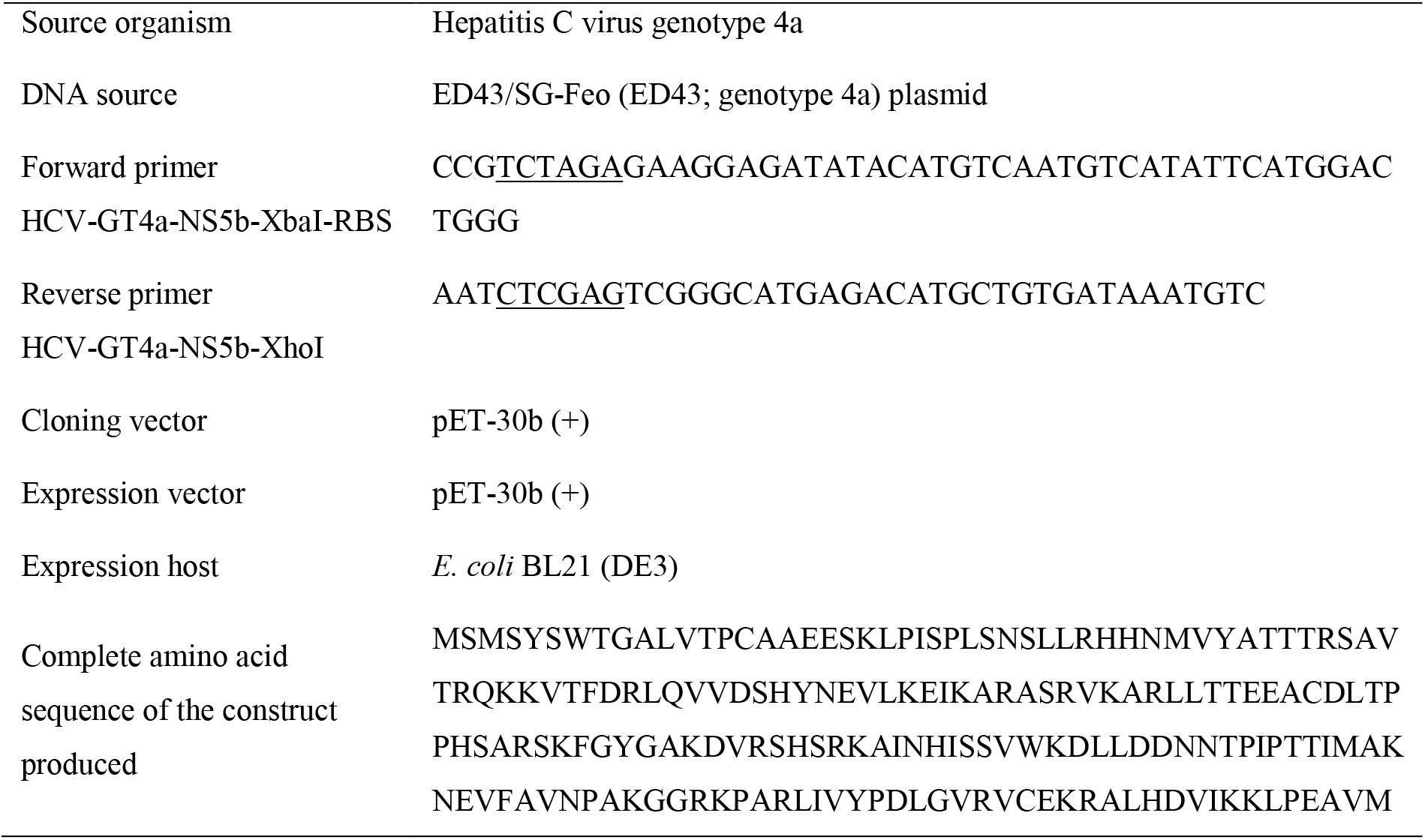
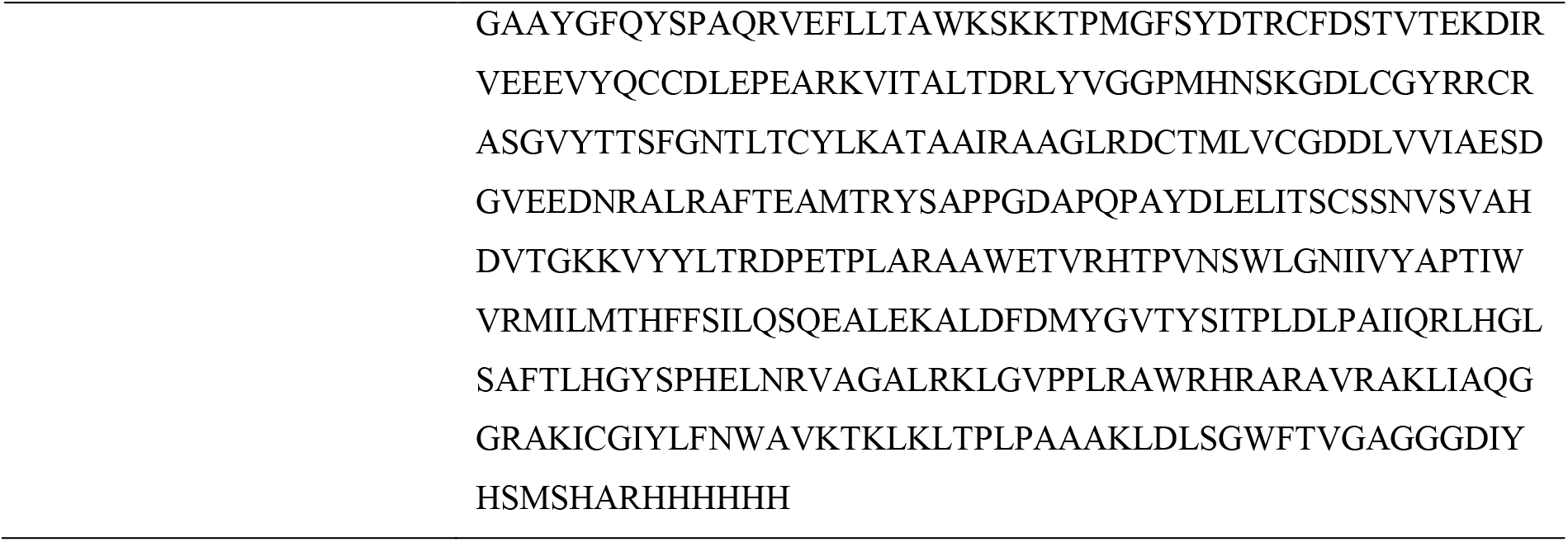
Macromolecule production information

**Table 2.** Crystallization of NS5b protein from hepatitis C virus genotype 4a.

|  |  |
| --- | --- |
| Method | Sitting-drop vapour diffusion |
| Plate type | Cryschem from Hampton Research |
| Temperature (K) | 292 |
| Protein concentration | 4 mg ml<sup>-1</sup> |
| Buffer composition of protein solution | 20 mM TRIS pH 7.5, 200 mM NaCl |
| Composition of reservoir solution | 100 mM sodium citrate pH 5.4, 25-30% (v/v) PEG 550 MME |
| Volume and ratio of drop | 3 µL, 1:1 |
| Volume of reservoir | 0.5 mL |

### 2.2. Crystallization

Crystallization experiments for NS5b were performed at the X-ray Crystallography Platform at Helmholtz Centre Munich. The initial crystallization screening of NS5b was set up at 292 K using 4 mg ml^-1^ of protein with a Mosquito (TTP Labtech) nanodrop dispenser in sitting-drop 96-well plates and commercial screens. The conditions were then optimized to obtain monocrystals of sufficient dimensions and quality for the X-ray diffraction experiments. Optimization was performed using the sitting-drop vapour diffusion method at 292 K in 24-well plates. The best diffracting crystals were obtained from the solution composed of 100 mM sodium citrate pH 5.4 and 25-30% (v/v) PEG 550 MME as a precipitating agent.

### 2.3. Data collection and processing

For the X-ray diffraction experiments, the NS5b crystals were mounted in a nylon loop and flash cooled to 100 K in liquid nitrogen. Diffraction data for HCV-NS5b crystals were collected on the ID30A-3 beamline (ESRF, Grenoble, France). Data collection was performed at 100 K. The dataset was indexed and integrated using XDS (Kabsch, 1993, 2010) and scaled using SCALA (Evans, 2006, Winn *et al*., 2011). Intensities were converted to structure-factor amplitudes using the program TRUNCATE (French & Wilson, 1978). Data collection and processing details are presented in Table 3.

**Table 3.** Data collection and processing Values for the outer shell are given in parentheses.

|  |  |
| --- | --- |
| Diffraction source | ID30A-3 beamline (ESRF, Grenoble, France) |
| Wavelength (Å) | 0.96500 |
| Temperature (K) | 100 |
| Detector | Pilatus3 2M |
| Crystal-detector distance (mm) | 310.14 |
| Rotation range per image (°) | 0.7 |
| Total rotation range (°) | 100 |
| Space group | $P2_12_12_1$ |
| $a, b, c$ (Å) | 63.3, 87.1, 96.7 |
| $\alpha, \beta, \gamma$ (°) | 90, 90, 90 |
| Mosaicity (°) | 0.32 |
| Resolution range (Å) | 50–3.10 (3.18–3.10) |
| Total No. of reflections | 39,278 |
| No. of unique reflections | 10,145 |
| Completeness (%) | 99.7 (100) |
| Redundancy | 3.9 (4.0) |
| $\langle I/\sigma(I) \rangle$ | 9.23 (1.74) |
| $R_{\text{mean}}$ | 13.8 (94.1) |
| CC(1/2) | 99.5 (71.3) |
| Overall $B$ factor from Wilson plot (Å<sup>2</sup>) | 77.2 |

### 2.4. Structure solution and refinement

The structure of NS5b genotype 4a was solved by molecular replacement with MolRep (Vagin & Teplyakov, 1997) from CCP4 (Winn *et al*., 2011) using the crystal structure of the RNA-dependent RNA polymerase from hepatitis C virus (PDB code: 1C2P, (Lesburg *et al*., 1999) as a search model. Both proteins share 75% sequence identity. Model rebuilding was done in COOT (Emsley *et al*., 2010). The refinement was done in REFMAC5 (Murshudov *et al*., 1997) using the maximum likelihood target function including TLS parameters (Winn *et al*., 2000). The final model is characterized by R_cryst_ and R_free_ factors of 20.2% and 25.7% (Table 4). The stereochemical analysis of the model was done in PROCHECK (Laskowski *et al*., 1993) and MolProbity (Chen *et al*., 2010). All software was part of the SBGrid software bundle (Morin *et al*., 2013). Comparison and analysis of the structure has been performed with Dali server (Holm *et al*., 2008).

**Table 4.**
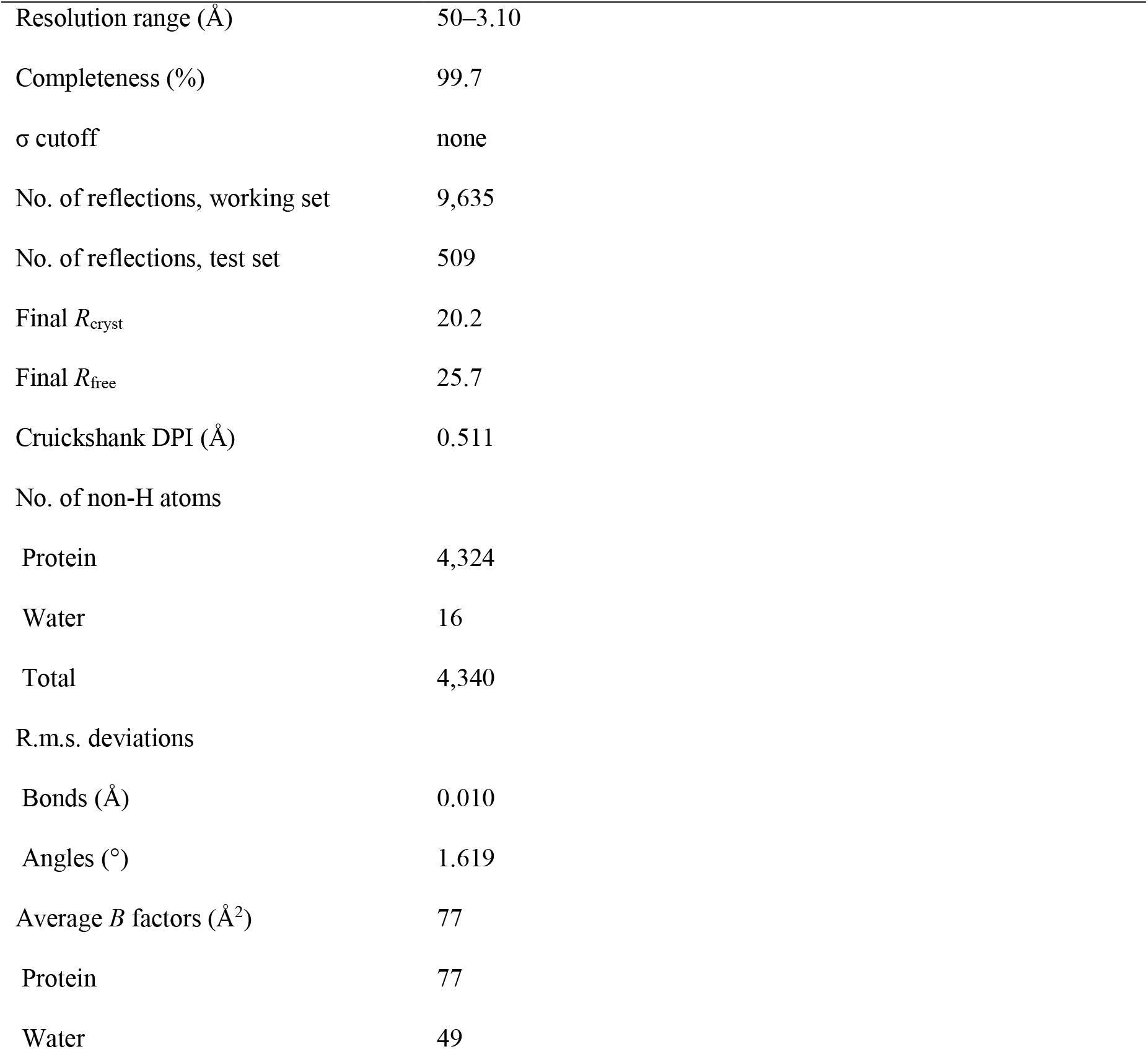
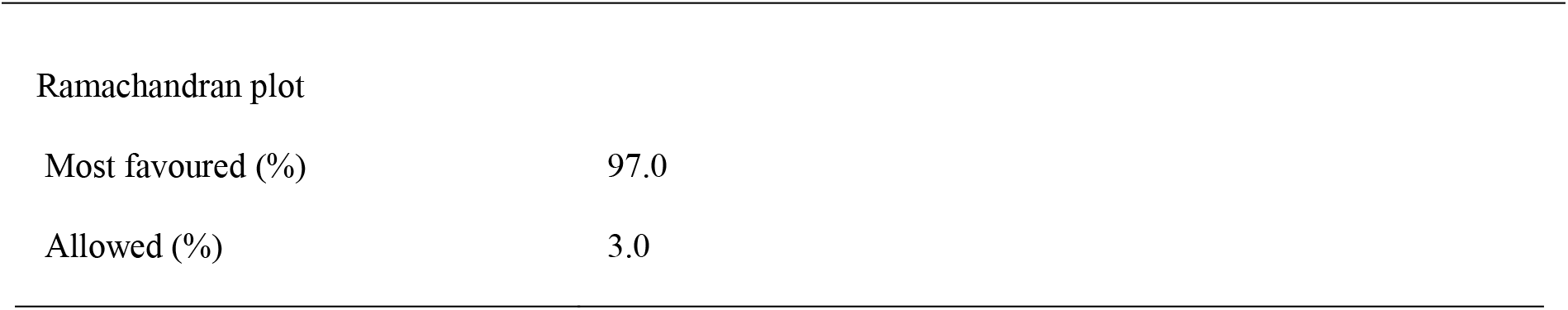
Structure solution and refinement

|  |  |
| --- | --- |
| Resolution range (Å) | 50–3.10 |
| Completeness (%) | 99.7 |
| $\sigma$ cutoff | none |
| No. of reflections, working set | 9,635 |
| No. of reflections, test set | 509 |
| Final $R_{\text{cryst}}$ | 20.2 |
| Final $R_{\text{free}}$ | 25.7 |
| Cruickshank DPI (Å) | 0.511 |
| No. of non-H atoms |  |
| Protein | 4,324 |
| Water | 16 |
| Total | 4,340 |
| R.m.s. deviations |  |
| Bonds (Å) | 0.010 |
| Angles (°) | 1.619 |
| Average $B$ factors (Å<sup>2</sup>) | 77 |
| Protein | 77 |
| Water | 49 |
| Ramachandran plot |  |
| Most favoured (%) | 97.0 |
| Allowed (%) | 3.0 |

## 3. Results and discussion

### Protein expression and purification

To solve the structure of NS5b protein from HCV genotype 4 by protein crystallization experiments, we cloned a full length as well as a truncated NS5b macromolecule into the bacterial expression vector pET-30b (+). Although we were able to successfully express and purify the C-terminally truncated NS5b protein, we failed to express a stable, full-length NS5b in bacterial cells. This observation is consistent with previous studies, which also reported difficulties in the expression of the NS5b full length in bacteria. One possible reason is the presence of a C-terminal membrane-anchor (Wang *et al*., 2004, Huang *et al*., 2004).

### Crystallization and structural determination

The initial crystallization screening of genotype 4 NS5b was set up at 292 K using 4 mg ml^-1^ of protein with a nano-drop dispenser in sitting-drop 96-well plates and commercial screens. Optimization was performed using the sitting-drop vapour diffusion method at 292 K in 24-well plates. Crystals of NS5b appeared after 2-3 days and required another few days to grow to the final size. The biggest crystals were obtained from PEG 550 MME as a precipitating agent. Although the morphology of the crystals was not ideal, we performed X-ray experiment and managed to collect satisfactory quality data set to 3.1 Å resolution. The structure has been successfully solved with the molecular replacement method, refined to the final R_cryst_ and R_free_ factors of 20.2% and 25.7%. Atomic coordinates and structure factors have been deposited in the Protein Data Bank under accession code 6gp9.

### Structure description

Similar to the available structures of other genotype HCV polymerases (Biswal *et al*., 2005, Powdrill *et al*., 2010, Karam *et al*., 2014), the three-dimensional fold of the HCV-NS5b genotype 4a protein could be depicted in the classical right-hand arrangement with fingers, palm, and thumb domains (Figure 1a). Structural comparison of the genotype 4a HCV-NS5b with other genotypes revealed HCV RNA polymerase genotype 2a as closest structural homolog (PDB ID: 1yv2; (Biswal *et al*., 2005)). Comparisons of both structures shows an root-mean-square (r.m.s.) deviation of 0.92 Å with a sequence identity between the two proteins of 74%. Figure 1b shows the overlay of both structures with a third HCV NS5b protein structure (PDB ID: 2wrm; unpublished) that shows the highest sequence identity of 79% to the protein studied here. Biswal *et al*. (2005) reported genotype 2a NS5b structure in an open, inactive (PDB ID: 1yv2) and closed, active (PDB ID: 1yuy) conformation. The r.m.s. deviation of the genotype 4 NS5b is 0.92 Å and 1.39 Å for the open and closed conformation, respectively. It clearly indicated that our final model adopts an open, inactive conformation, which was expected for our crystallization experiments, as we did not use any NS5b substrates or RNA template.

**Figure 1.**
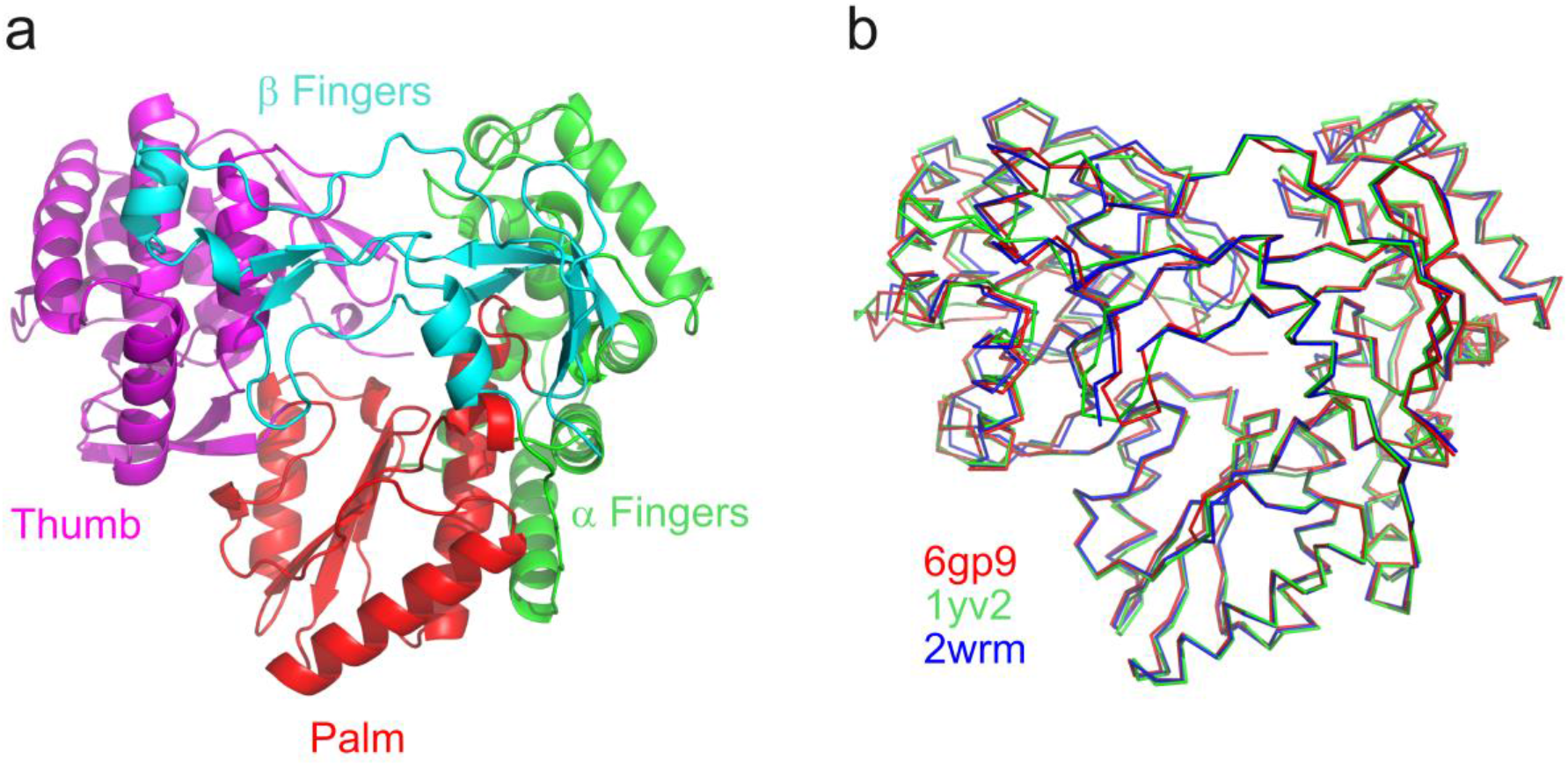
**a)** The overall structure of genotype 4 HCV NS5b protein shown as a cartoon with domains presented in different color: thumb – magenta, palm – red, α fingers – green, β fingers - cyan (color code as in Ago *et al*., 1999). **b)** Superposition of genotype 4 HCV NS5b protein (PDB ID: 6gp9 shown as red ribbon) with the closest structural homologue, HCV RNA polymerase genotype 2a (Biswal *et al*. 2005; PDB ID: 1yv2; shown in green), and the HCV RNA polymerase showing the highest sequence identity (PDB ID: 2wrm; unpublished; shown in navy blue).

Taken together, in this research paper we have solved the crystal structure of the HCV-NS5b RNA polymerase of the Egyptian genotype 4a. Structural similarity to NS5b of other HCV genotypes (Figure 1b), as well as conservation of the active site (unpublished data) indicate that NS5b inhibitors are very likely to act in a true pan-genotypic fashion.

## Acknowledgements

We thank Prof. Dr. Charles Rice, New York, for kindly providing the ED43/SG-Feo (ED43; genotype 4a) plasmid. We thank Vera Roman, Helmholtz Zentrum München, for her kind support.

## Supporting information

### S1. Western blot analysis of HCV NS5b

To confirm the expression of His-Tagged HCV NS5b, proteins were resolved by SDS-PAGE and after electrophoresis transferred to a polyvinylidene difluoride membrane (PVDF) (GE, health) using a Trans-Blot SD Semi-Dry Transfer Cell (Bio-Rad, Munich, Germany). PVDF membrane was activated by soaking into methanol for 1 min. Then all components were soaked in 1X transfer buffer and the blotting sandwich was layered as follows: from bottom to top: sponge layered with filter paper followed by the gel, the PVDF membrane, a piece of filter paper and a pad. The assembly was then run at a constant run at 18V for an hour. The membrane was then blocked for non-specific binding by incubation in blocking buffer (TBS-T containing 5% milk powder). The membrane was then incubated at 4°C in m anti-His primary antibody diluted in (TBS-T containing 5% milk powder) at a dilution of (1:2000). The membrane was then washed 3X 10 min with (TBST), then the membrane was incubated with HRP-secondary antibody at a concentration of 1:10,000 for one hour at room temperature. The membrane was then washed three times as described above. Bound antibodies were detected by ECL+-system (Amersham) and protein bands were visualized by Fusion.

**Figure S1.**
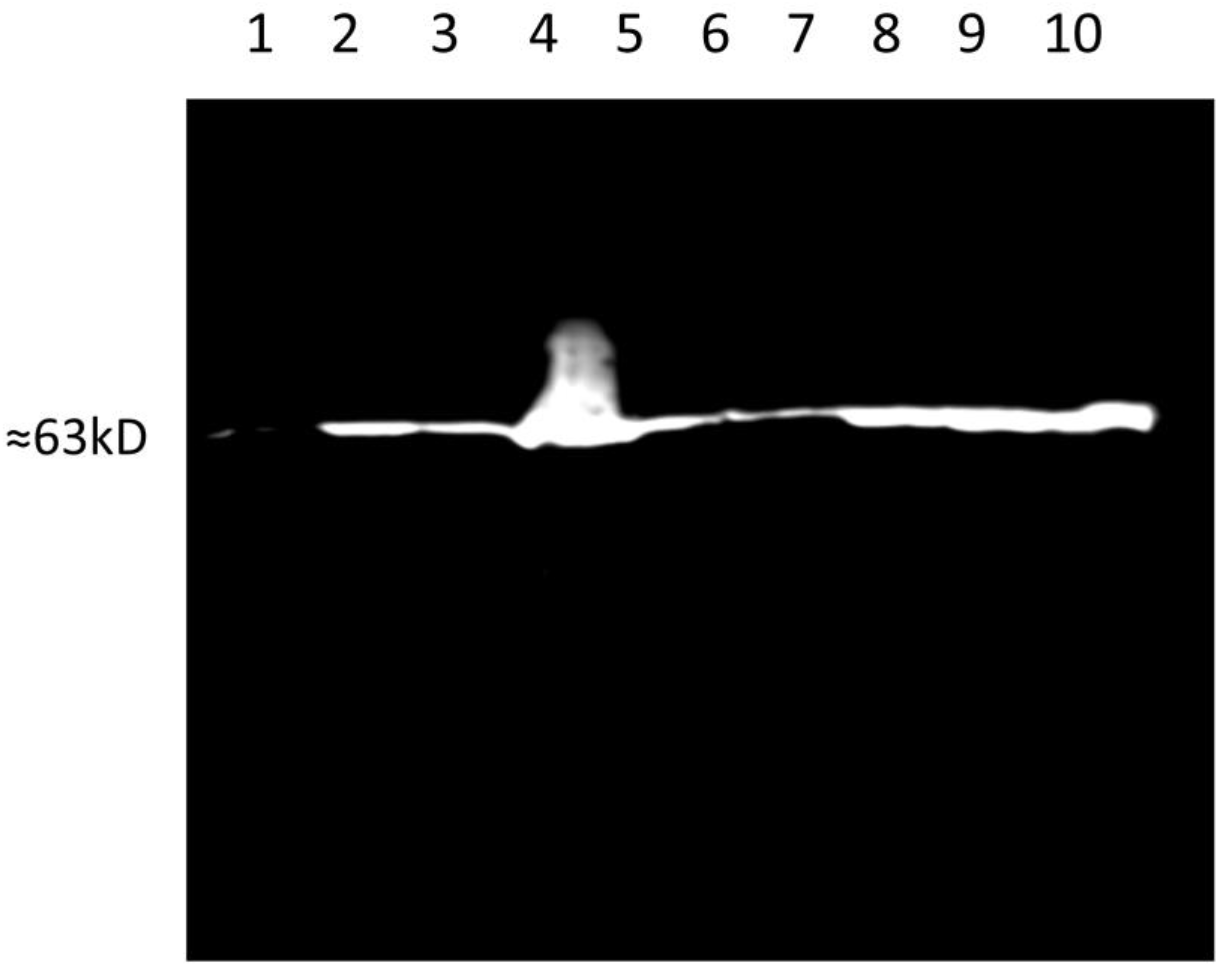
Western blot analysis of NS5BΔ23-His6: Lane 1 is lysate before protein induction, lane 2 is lysate after induction, lane 3 is lysate cleared by centrifugation, lane 4 is pellet, lane 5 is the first wash, lane 6 is the second wash, lane 7 is the first elution, lane 8 is the second elution, lane 9 is the third elution and lane 10 is flow-through.

